# Unravelling the molecular mechanisms of DNA capture by the Com pilus in naturally transformable monoderm bacteria

**DOI:** 10.1101/2025.03.12.642782

**Authors:** Jérémy Mom, Odile Valette, Laetitia Pieulle, Vladimir Pelicic

## Abstract

Transformation is a mechanism of horizontal gene transfer widespread in bacteria. The first step in transformation – capture of exogenous DNA – is mediated by surface-exposed filaments belonging to the type 4 filament (T4F) superfamily. How these protein polymers, composed of major and minor pilin subunits, interact with DNA remains poorly understood. Here, we address this question for the Com pilus, a widespread T4F mediating DNA capture in competent monoderm species. Our functional analysis, performed in *Streptococcus sanguinis*, was guided by a complete structural model of the Com pilus. We show that the major pilin ComGC does not bind DNA. In contrast, a systematic mutational analysis of electropositive residues exposed at the filament surface in the four minor pilins (ComGD, ComGE, ComGF, ComGG) reveals that the interface between ComGD and ComGF is important for DNA capture. Sequential mutations in these two interacting subunits lead to complete abolition of transformation, without affecting piliation. We further demonstrate the physical interaction between ComGD and ComGF using disulfide crosslinking, upon mutagenesis of two strategically positioned residues into cysteines. A structural model of the Com pilus tip interacting with DNA recapitulates all these findings and highlights a novel mode of DNA-binding, conserved in hundreds of monoderm species.

**IMPORTANCE:** Bacteria are capable of evolving and diversifying very rapidly by acquiring new genetic material via horizontal gene transfer (HGT). Transformation is a widespread mechanism of HGT in bacteria, which results from the capture of extracellular DNA by surface-exposed pili belonging to the superfamily of type 4 filament (T4F). How T4F – that are composed of major and minor pilins – interact with DNA remains poorly understood, especially in competent monoderm species that use a unique T4F for DNA capture known as Com pilus or T4dP. The significance of this work is in characterizing a novel mode of DNA-binding by showing that the interface between two minor pilins part of a tip-located complex of four pilins – found in many different T4F – have been functionalized in monoderms to capture DNA. This is an evolutionary strategy used by bacteria to promote the exceptional functional versatility of T4F.

## INTRODUCTION

Bacteria are capable of evolving and diversifying very rapidly, which has allowed them to colonize and thrive even in the most extreme environments. Since they replicate clonally by binary fission, bacterial extraordinary evolutionary potential results from their ability to acquire new genetic material by horizontal gene transfer (HGT) (1). Deciphering the mechanisms of HGT in bacteria – that occurs by transformation, transduction, or conjugation – has been a hot topic for research for decades, also because HGT has dire consequences for human health by leading to the spread of antibiotic resistance and the evolution of more virulent strains (1).

Transformation – documented in a wide variety of bacterial species (2) – is the only mode of HGT that depends exclusively on the recipient cell, which must be in a genetically programmed state called competence (3). Transformation is defined as a genetic alteration of the recipient resulting from (i) the capture of free DNA from the extracellular milieu (also known as DNA uptake), (ii) its translocation across the cytoplasmic membrane (CM), and (iii) its stable integration into the genome (3). At a molecular level, DNA capture results from extracellular DNA being bound by filaments on the cell surface (4), translocated across the outer membrane and/or the cell wall following filament retraction (4), and compacted in the vicinity of the CM upon interaction with the DNA receptor ComEA (5, 6). Then, a single strand of DNA is translocated across the CM through the ComEC channel (7), while the second strand is degraded. Finally, internalized DNA is integrated in the genome by RecA-mediated strand exchange.

With only one known exception, binding of extracellular DNA – the earliest step in transformation – requires one of the several subtypes of type 4 pili (T4P) (3). T4P belong to the superfamily of type 4 filaments (T4F) – functionally versatile nanomachines ubiquitous in Bacteria and Archaea (8, 9) – that use a conserved multi-protein machinery to assemble and operate a filamentous polymer composed of type 4 pilins. Pilins are defined as major or minor according to their respective abundance in filaments (10). Although we still lack a detailed mechanistic understanding of DNA capture by T4P, the following scenario is widely admitted. As visualized in competent diderm species – that all use type 4a pili (T4aP) to capture DNA, where “a” denotes the subtype – filaments bind DNA directly (4, 11). A few pilins acting as DNA receptors have been identified, of which the best characterized are *Neisseria meningitidis* ComP (11) and *Legionella pneumophila* FimT (12). These unrelated pilins bind DNA via patches of electropositive residues exposed on the filaments surface. ComP binds DNA via an electropositive stripe delimited by two disulfide bonds (13), and shows a preference for a short sequence motif repeated hundreds of times in *Neisseria* genomes, explaining these species uncommon preferential uptake of their own DNA (14). FimT binds DNA via a patch of conserved electropositive residues, located in a flexible/unstructured C-terminal tail (12). Subsequently, bound DNA is brought to the CM by T4aP retraction powered by the retraction motor PilT (15). Preventing pilus retraction, either by *pilT* mutagenesis (16) or by steric obstruction (4), prevents DNA uptake.

The above DNA-binding pilins are found only in a small subset of competent species, which suggests that other modes of DNA capture exist. In particular, no ComP or FimT homologs are present in T4dP – also known as Com pili (17) – that are found in hundreds of competent monoderm species (9). T4dP, which are long (18) and dynamic filaments (19), represent a minimalistic T4F (17) requiring only a set of proteins universally conserved in the T4F superfamily (20) – pilins, prepilin peptidase (PPase), extension ATPase and platform protein – for filament assembly and functioning (17). Here, using *Streptococcus sanguinis* as a model, we determined how T4dP bind DNA to promote its capture. We performed a functional analysis guided by a complete structural model of the filament composed of five pilins (ComGC, ComGD, ComGE, ComGF, ComGG), which uncovered a novel mode of DNA-binding involving two adjacent tip-located pilins. We show that the resulting model is applicable to hundreds of monoderm species that express T4dP.

## RESULTS

### T4dP display a canonical T4P structure with a filament capped by a complex of four minor pilins

Recently, using *S. sanguinis* as a model, we showed that the eight *com* genes organized in two transcription units (*comGA*-*comGB*-*comGC*-*comGD*-*comGE*-*comGF*-*comGG* and *comC*) (Fig. S1A) – conserved in hundreds of species of Firmicutes (9) – are necessary and sufficient for the assembly and functioning of T4dP (17). The identification of signature domains in the corresponding proteins using InterProScan (21), and the prediction of their 3D structures using AlphaFold 2 (22) showed that they correspond to the four proteins universally conserved in T4F (17): PPase (ComC), extension ATPase (ComGA), platform protein (ComGB), and type 4 pilins (ComGC, ComGD, ComGE, ComGF and ComGG). We confirmed that the last five proteins are processed by the PPase ComC – *i.e.*, their N-terminal class 3 signal peptide (SP3) found in all type 4 pilins (10) is cleaved by ComC (Fig. S1B) – and assembled into filaments (17). ComGC is the major pilin, while the remaining four (ComGD, ComGE, ComGF and ComGG) are minor pilins much less abundant in pilus preparations. We showed that the four minor pilins interact to form a complex (17), predicted to be located at the tip of T4dP based on similarities to complexes of four minor pilins found at the tip of other better characterized T4F (23, 24).

As a first step to understand how T4dP bind DNA to promote its capture during transformation, we modelled the filament by taking advantage of the recent introduction of AlphaFold 3, a deep-learning framework with substantially improved prediction accuracy for complex structures (25). Using the sequence of the mature pilins of *S. sanguinis*, in which the SP3 cleaved by ComC (Fig. S1B) were manually removed, we produced a complete structural model of T4dP. As described previously, when modelled on their own, each pilin displays a “lollipop” architecture typical of type 4 pilins (10), starting with an α-helix (α1) of approximately 50 residues onto which a globular head is impaled (Fig. S2). The hydrophobic N-terminal half of α1 (α1N) protrudes from the pilin subunits and packs within the filament core upon pilus polymerization. All the five pilin structural models exhibit excellent confidence metrics (Fig. S2), except for the last 30 residues of ComGG for which the local confidence is very low. This portion of ComGG, although it is modelled as an α-helix, protrudes at an “unnatural” angle from the globular head of this pilin (Fig. S2). Next, we produced a complete model of a T4dP composed of 10 ComGC subunits and one subunit each of the four minor pilins (Fig. 1). Critically, this was not feasible using AlphaFold 2, which only allowed us to model a complex of four minor pilins (17). The complete model reveals that T4dP display a canonical T4P structure (Fig. 1A). The ComGC pilus shaft is similar to available near-atomic resolution T4aP structures (26–28). The major pilins pack helically with their α1 helices forming a hydrophobic core, which is accompanied by a “melting” of a portion of α1N that becomes non-helical (Fig. 1A). Strikingly, the filament is capped by a ComGD-ComGF-ComGE-ComGG complex of four minor pilins (Fig. 1B), which were previously shown to interact in *S. sanguinis* (17). As for ComGC, packing of ComGD is accompanied by partial melting of α1N (Fig. 1B). Considering its size, the confidence metrics for the T4dP structural model are good. The quality of the model, estimated using the structure assessment tool in SWISS-MODEL (29), returned a MolProbity score of 2.43 and a global QMEANDisCo score of 0.67 ± 0.05. These values are comparable to those calculated for the recently determined cryo-EM structure of *S. sanguinis* T4aP (28) (MolProbity 1.12 and global QMEANDisCo 0.65 ± 0.05). Intriguingly, the local confidence for the last 30 residues of ComGG remained very low and the corresponding α-helix still protrudes unnaturally from the tip of the pilus (Fig. 1).

**Fig. 1.**
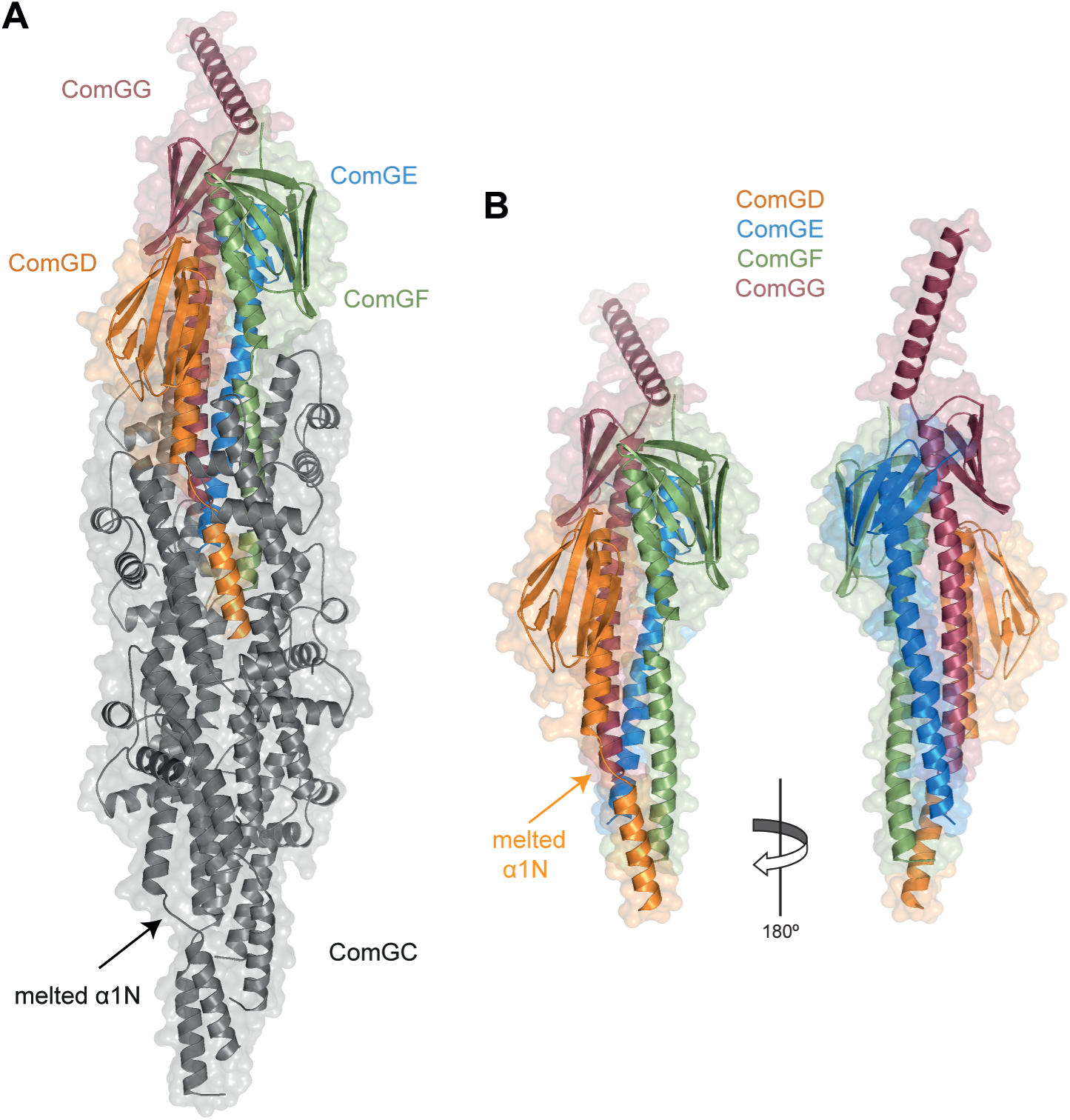
Modelling of T4dP reveals a canonical T4P, with a ComGC filament capped by a complex of four minor pilins. **A**) Structural T4dP model in *S. sanguinis* predicted by AlphaFold 3 (25) with 10 copies of ComGC and one copy of each the four minor pilins. The cartoon representation with surfaces shown in transparency reveals that T4dP display a canonical T4P structure, where a filament of ComGC (grey) is capped by a complex of four minor pilins ComGD (orange), ComGE (blue), ComGE (green), ComGG (maroon). The same color code is used throughout the manuscript. As in other available near-atomic resolution structures of T4aP (26–28), helical packing of ComGC subunits via their α1 helices is accompanied by “melting” of a portion of α1N that becomes non-helical. **B**) Focus on the tip-located complex of ComGD-ComGF-ComGE-ComGG minor pilins. Two different sides are shown (180° views). As for ComGC, packing of ComGD in the pilus is accompanied by partial melting of its α1N. Intriguingly, the local confidence for the last 30 residues of ComGG, which was very low when this pilin was modelled on its own (see Fig. S2), remained very low and the corresponding α-helix still protrudes unnaturally from the tip of the pilus.

Our modelling results strengthen the notion that T4dP are canonical T4P where a ComGC pilus shaft is capped by a complex of four conserved minor pilins. This architecture is similar to that of better characterized T4F.

### ComGC has no intrinsic DNA-binding propensity

In order to answer the question which subunit of the T4dP binds DNA to promote its capture during transformation, we first focused on ComGC. To determine whether this major pilin has intrinsic DNA-binding activity, we tested its propensity to interact with DNA using agarose electrophoretic mobility shift assays (EMSA), as previously done for ComP from *Neisseria* (11). We used the soluble portion of ComGC – the globular head without the hydrophobic α1N, which was previously characterized structurally (30) – fused to maltose-binding protein (MBP) for purification. As a target probe, we used the pUC19 plasmid. As a positive control, we used an MBP fusion to *S. sanguinis* ComEA, the DNA receptor involved in the late stages of DNA uptake (5, 6). Since ComEA is a bimodular protein in *S. sanguinis*, as revealed by InterProScan (21) (Fig. S3A), we fused only the C-terminal ComEA module (IPR004509) to MBP. This module is structurally similar to the ComEA crystal structure of *Thermus thermophilus* (Fig. S3B), which consists of a repeated helix-hairpin-helix region. The EMSA showed that ComGC is not capable of binding DNA since no shift was seen with as much as 10 µM of purified protein (Fig. 2). In contrast, in the same conditions, ComEA from *S. sanguinis* bound DNA efficiently (Fig. 2). We observed a shift with as little as 0.5 µM protein, indicating a significant affinity of *S. sanguinis* ComEA for DNA, as shown in other species (5, 6). These experiments rule out the possibility that ComGC, the major subunit of T4dP, might be the primary DNA receptor during DNA capture. This suggests that the four minor pilins that cap the pilus are likely be involved in DNA-binding to initiate transformation.

**Fig. 2.**
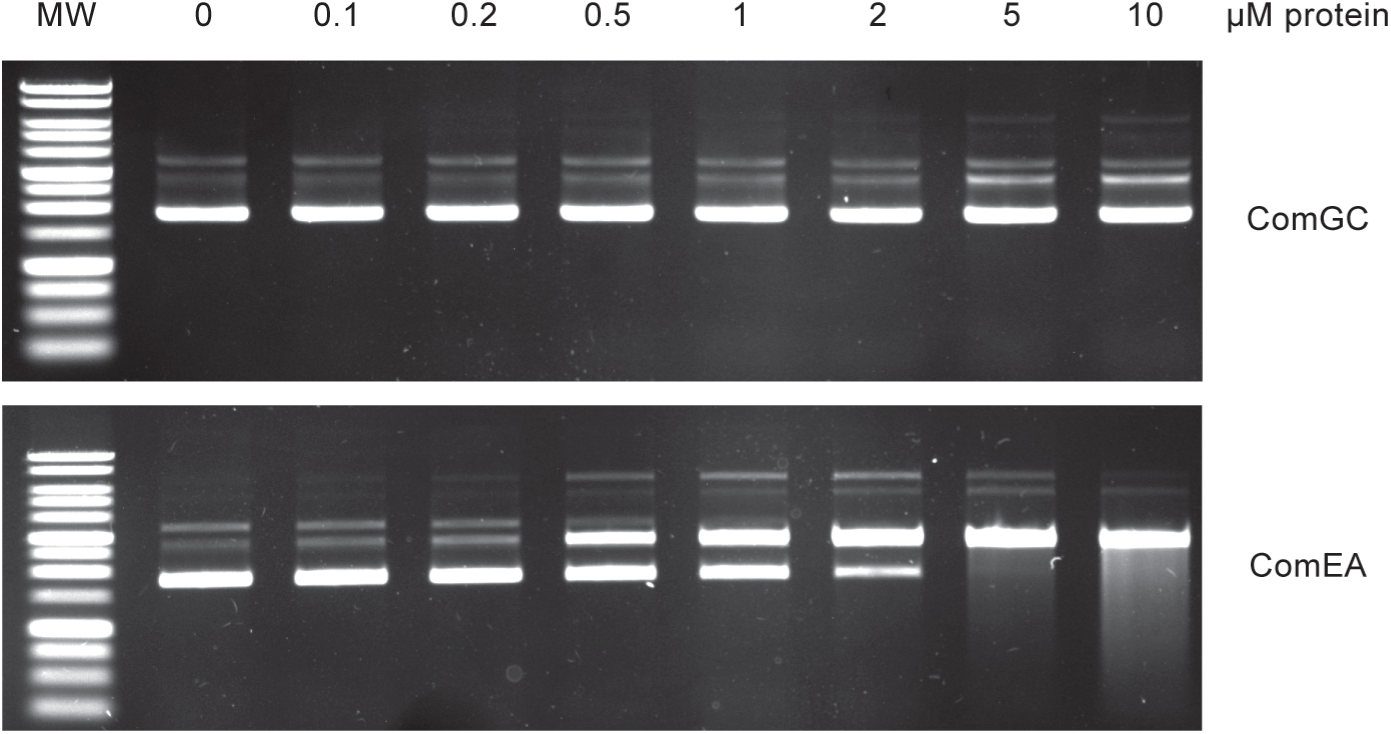
ComGC has no intrinsic DNA-binding activity. The DNA-binding propensity of purified MBP-ComGC was assessed by agarose EMSA. A standard amount (120 ng) of pUC19 plasmid was incubated with increasing concentrations of purified MBP-ComGC and resolved by electrophoresis on a 0.8 % agarose gel (upper panel). As a positive control, we used purified MBP-ComEA (lower panel). ComEA is a conserved DNA receptor involved in the late stages of DNA uptake (5).

### Surface-exposed electropositive residues in ComGD and ComGF are key for DNA capture by Com pili

We next focused on the four minor pilins ComGD, ComGE, ComGF and ComGG to understand how the pilus binds DNA to promote its capture during transformation. Since these proteins were recalcitrant to purification despite multiple attempts, we could not determine whether they exhibit intrinsic DNA-binding activity by EMSA. Instead, guided by our structural model, we made a series of mutations in each of the four minor pilin-encoding genes in *S. sanguinis* and tested their effects on transformation as a proxy for DNA binding. When phenotypic defects in transformation were observed, we ruled out that these could result from impaired pilus biogenesis by testing the mutants for piliation. We used a previously designed pilus purification procedure (17), and detected the pili by immunoblotting using an anti-ComGC antibody.

Since FimT was reported to bind DNA via an electropositive flexible/unstructured C-terminal tail (12), we first focused on the C-terminal tail in ComGG (Fig. 3A), which is invariably modelled with very poor confidence. As seen by modelling other ComGG (Fig. S4A), this structural feature is widely conserved in *Streptococcus*-type ComGG (IPR047665) that are found mainly in *Streptococcaceae* and *Enterococcaceae*. As seen by aligning the sequences of the ComGG tails in *S. sanguinis* proteins available in InterPro (Fig. 3B), 20 out of 30 residues in this region are charged, ten of which are positively charged Lys. This sequence feature is conserved in other ComGG, such as in *S. pneumoniae* proteins (Fig. S4B). We therefore engineered a mutant in *S. sanguinis* expressing a shorter ComGG, ComGG_Δ94-122_, in which the tail residues were truncated. The unmarked mutant has a stop codon in *comGG* instead of the codon encoding Lys_94_ (the numbering is according to the processed protein). Since pilus production was previously reported to be abolished in a *ΔcomGG* mutant (17), we first determined whether the *comGG_Δ94-122_* mutant is piliated by analyzing pilus preparations by immunoblotting using an antibody against the major pilin ComGC. Critically, we found that the *comGG_Δ94-122_* mutant is unaffected for piliation despite the deletion of almost a quarter of the ComGG protein (Fig. 3C). Next, we tested whether this mutant was still transformable by quantifying its competence. We found that the *comGG_Δ94-122_* mutant was as transformable as the wild-type (WT) strain (Fig. 3D), *i.e.*, 4.84 ± 1.79 % of transformed cells compared to 5.69 ± 1.43 %, respectively. In contrast, we previously showed that transformation was abolished in non-piliated mutants, with decreases at least 6 orders in magnitude (17). Taken together, these findings demonstrate that the charged C-terminal tail in ComGG is dispensable for DNA binding and uptake by T4dP.

**Fig. 3.**
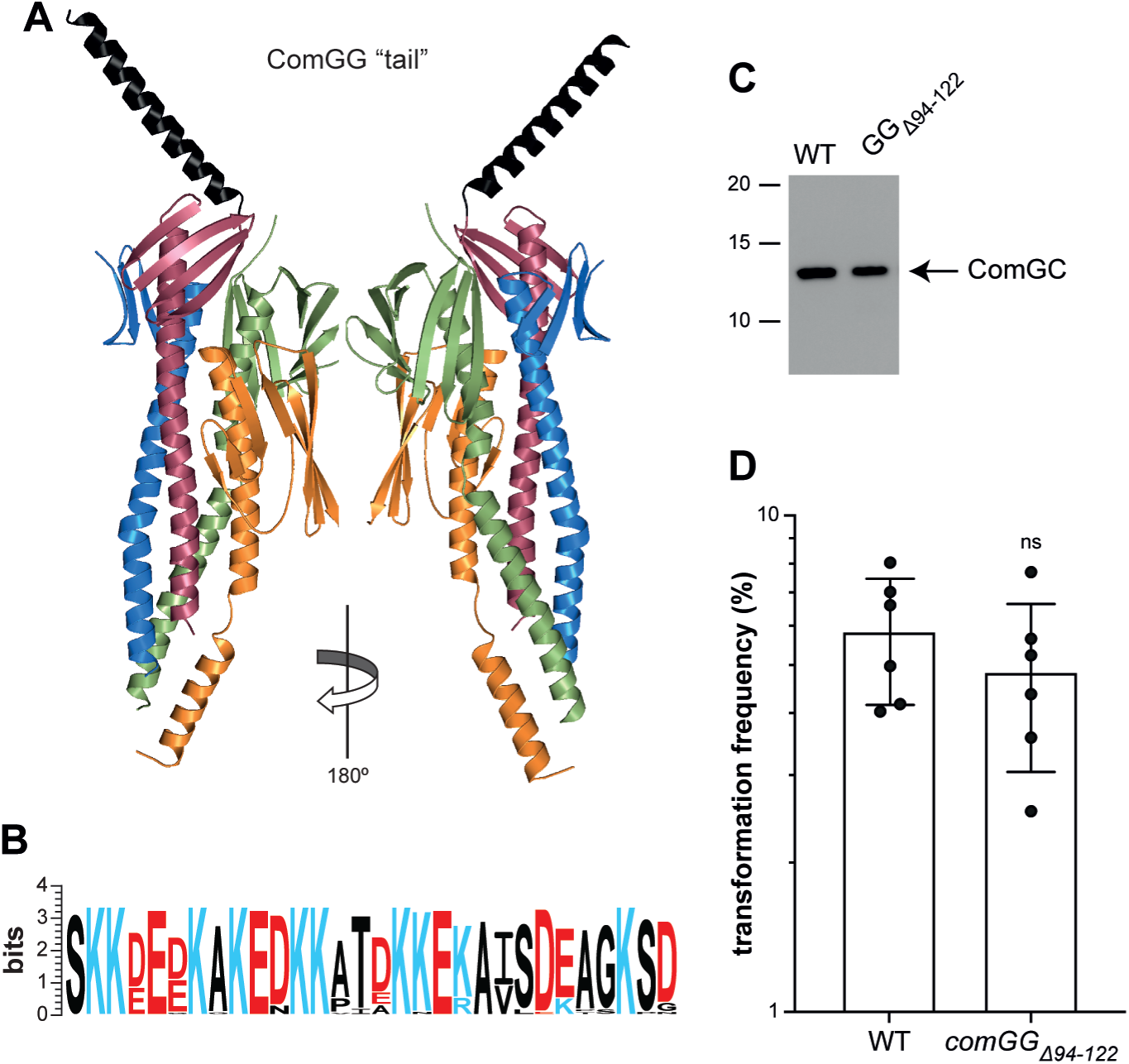
The charged C-terminal tail in ComGG is dispensable for *S. sanguinis* piliation and transformation. **A**) Tip-located complex of four minor pilins with the last 30 residues of ComGG, which form a α-helical tail with very low local modelling confidence, highlighted in black. Two different sides are shown (180° views). **B**) The ComGG tail is highly charged as could be seen from the sequence logo generated from a MSA of 31 *S. sanguinis* IPR047665 entries (*Streptococcus*-type ComGG) in InterPro (43). Charged residues are colored in blue (electropositive) or red (electronegative). **C**) An unmarked tail-less *S. sanguinis* mutant expressing ComGG_Δ94-122_ is piliated. Piliation was assessed by immunoblotting using an anti-ComGC antibody on pilus preparations made from equal volumes of culture. The WT strain is included as a control. **D**) A tail-less *S. sanguinis comGG_Δ94-122_* mutant is transformable. The WT strain is included as a control. Transformation frequencies (%) – mean ± SD from six independent experiments – are the ratio of transformants relative to number of viable bacteria. The *comGG_Δ94-122_* mutant is as transformable as the WT strain as assessed by a two-tailed t test. ns, not statistically different.

Next, because the *Neisseria*-specific minor pilin ComP was reported to bind DNA via a surface-exposed stripe of electropositive residues (11), we focused on the electropositive residues in the four minor pilins predicted to be exposed on the surface of T4dP. We systematically constructed a series of *S. sanguinis* mutants expressing minor pilins in which all positively charged and surface-exposed Lys and Arg residues were substituted into Gln. We started with eight mutations in ComGD and found that three mutants – K_101_Q, K_121_Q, K_123_Q – are significantly less competent, with 10- to 100-fold decreases in transformation (Fig. 4A). Strikingly, these residues were spatially close on ComGD, suggesting that they together form a DNA-binding site, which was supported by combining the mutations. Indeed, ComGD polymutants – K_121_Q/K_123_Q and K_101_Q/K_121_Q/K_123_Q – displayed even more drastic defects in transformation (Fig. 4A), with up to a 1,000-fold decrease. Importantly, all these mutants were still piliated as demonstrated by our ability to purify and detect T4dP (Fig. 4E). In contrast, the four mutants that we constructed in ComGE were as competent for transformation as the WT strain, suggesting that this minor pilin has no role in DNA binding (Fig. 4B). For ComGF, we obtained similar results than for ComGD. Four of the 10 mutants we constructed – K_65_Q, R_73_Q, K_81_Q, R_93_Q – were significantly less competent than WT (Fig. 4C), with up to a 1,000-fold decrease in transformation. As for ComGD, these residues were spatially close on ComGF, suggesting that they form a DNA-binding site. This was strengthened by showing that the R_73_Q/R_93_Q polymutant displayed an even more drastic, 10^4^-fold, decrease in transformation (Fig. 4C). Importantly, all these mutants were still piliated as demonstrated by our ability to purify and detect T4dP (Fig. 4E). Finally, For ComGG, we obtained similar results than for ComGE. None of the four mutants that we constructed in ComGG – we did not target the electropositive residues in the tail that were demonstrated in the experiments above to play no role in DNA binding (see Fig. 3) – were affected for competence (Fig. 4D), suggesting that this minor pilin has no role in DNA binding.

**Fig. 4.**
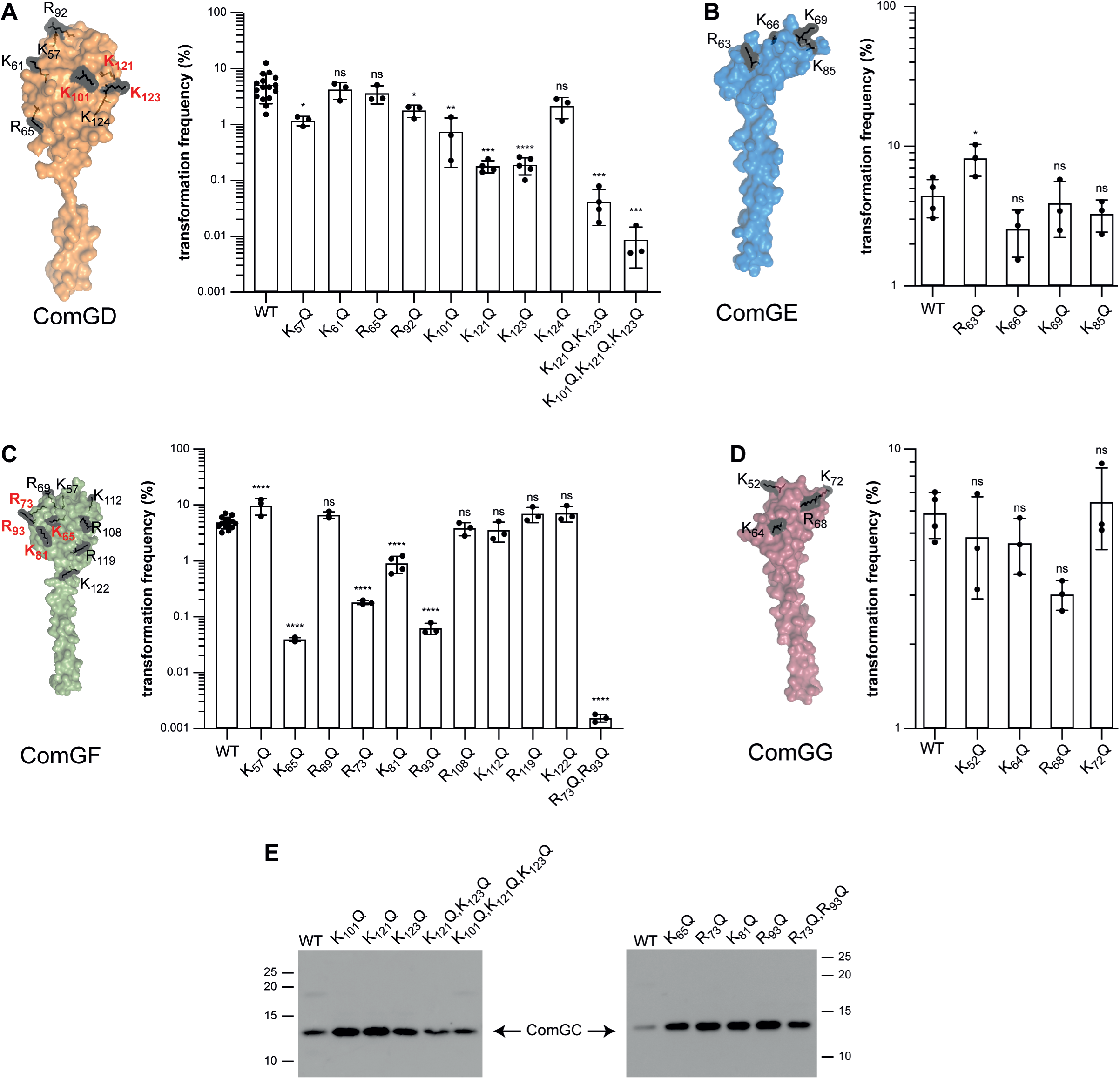
Electropositive residues in ComGD and ComGF exposed on the filament surface are key for transformation. **A**-**D**) Quantifying transformation in *S. sanguinis* mutants expressing minor pilins in which we altered surface-exposed electropositive residues potentially contributing to DNA binding. The targeted residues are highlighted in black on the corresponding structures in surface representation (the C-terminal tail in ComGG is not shown). Transformation frequencies (%) were the mean ± SD from at least three independent experiments. We used Dunnett’s one-way ANOVA to compare the means to the WT. ns, not statistically different; *, *P* 0.0332; **, *P* 0.0021; ***, *P* 0.0002; ****, *P* < 0.0001. **E**) Assessing piliation in the mutants affected for transformation. This was done by immunoblotting using an anti-ComGC antibody on pilus preparations made from equal volumes of culture. The WT strain is included as a control.

Taken together, these findings demonstrate that electropositive residues in the ComGD and ComGF subunits of the complex of four minor pilins capping T4dP are key for competence, most likely by playing a role in binding DNA.

### The interface between ComGD and ComGF at the pilus tip is key for DNA capture by T4dP

In the structural model of the pilus, ComGD and ComGF are adjacent subunits. Strikingly, the electropositive residues in these two minor pilins which are key for DNA capture, are spatially close on one side of the pilus tip complex (Fig. 5A), and could together represent an electropositive patch involved in binding DNA. To provide evidence for this, we combined mutations in ComGD and ComGF to create the quintuple mutant ComGD_K101Q/K121Q/K123Q_ ComGF_R73Q/R93Q_. Therefore, this mutant, which we will refer to as the 5Q mutant for the sake of readability, harbors as many as five substitutions of electropositive residues in two different minor pilins. We first showed that the 5Q mutant is piliated since ComGC could be detected by immunoblotting in pilus preparations of the 5Q mutant (Fig. 5B). Moreover, as shown by transmission electron microscopy (TEM), purified filaments displayed a morphology indistinguishable from WT filaments (17) (Fig. 5C). Critically, although the pili in the 5Q mutant appear normal, its transformation is abolished with a more than a 10^6^-fold decrease compared to WT (Fig. 5D). Such a dramatic defect in transformation, which has previously been observed only in non-piliated mutants (17), is stronger than the decreases observed in either of the polymutants in ComGD (10^3^-fold) or ComGF (10^4^-fold) (see Fig. 4). Together, these results show that electropositive residues in ComGD and ComGF on one side of the pilus tip constitute together an interface key for DNA capture by T4dP.

**Fig. 5.**
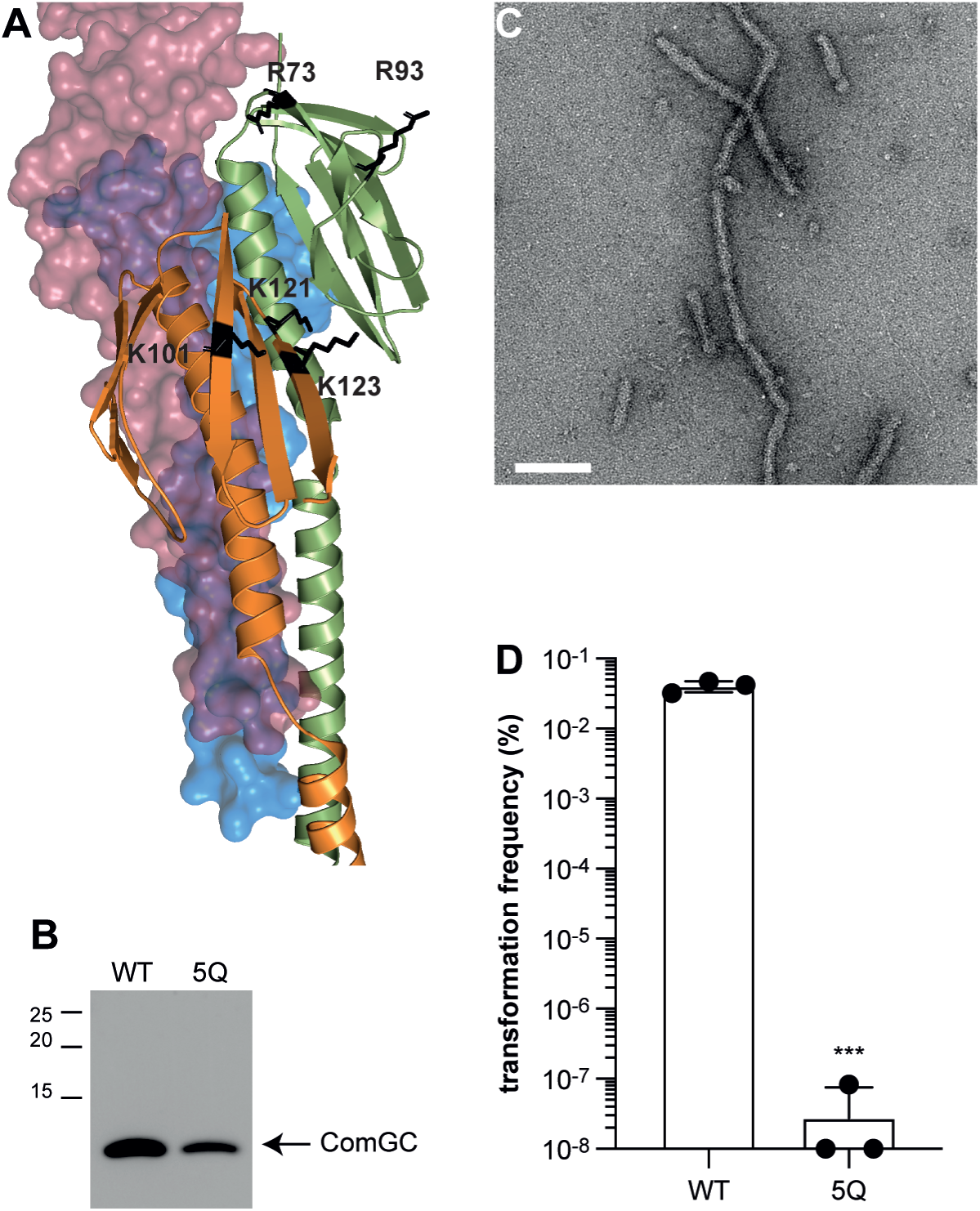
The interface between ComGD and ComGF is key for DNA capture by T4dP. **A**) ComGD-ComGF interface at the tip of the pilus. The electropositive residues shown to be important for transformation in Fig. 4 are highlighted in black. We constructed a quintuple *S. sanguinis* ComGD_K101Q/K121Q/K123Q_ ComGF_R73Q/R93Q_ mutant – named 5Q – in which all these electropositive residues were altered simultaneously. **B**) The 5Q mutant is piliated as assessed by immunoblotting using an anti-ComGC antibody on pilus preparations made from equal volumes of culture. The WT strain is included as a control. **C**) Pili in the 5Q mutant are morphologically normal as assessed by TEM on purified filaments. The scale bar represents 100 nm. **D**) Transformation is abolished in the 5Q mutant. Transformation frequencies (%) are the mean ± SD from three independent experiments – are the ratio of transformants relative to number of viable bacteria. Statistical significance assessed by a two-tailed t test. *** 0.0021 < *P* < 0.0002.

Next, to strengthen these findings, we confirmed the validity of the structural model by experimentally probing the predicted ComGD-ComGF interface in the context of the pilus. To do this, we used an *in vivo* disulfide crosslinking approach. (31). Guided by our structural model, we chose two pairs of residues in ComGD (Leu_52_ and Gly_118_) and ComGF (Asp_47_ and Gln_96_), which fall within disulfide crosslinking distance when the two proteins interact in the pilus. To facilitate detection of minor pilins, we constructed *S. sanguinis* single and double mutants in a strain that constitutively expresses T4dP (17), with Cys substitutions in the pairs of residues ComGD_L52C_/ComGF_D47C_ (Fig. 6A) and ComGD_G118C_/ComGF_Q96C_ (Fig. 6B). After pilus purifications, filaments were briefly treated with the thiol-specific oxidizer 4,40-dipyridyl disulfide (4-DPS) to promote the formation of disulfide bonds (32). Using anti-ComGD and anti-ComGF antibodies, we detected disulfide-bonded ComGD-ComGF adducts by immunoblotting after non-reducing gel electrophoresis in each of the two double mutants, but not in the respective single mutants (Fig. 6). Critically, in reducing electrophoresis conditions – in the presence of β-mercaptoethanol (β-ME) – the disulfide-bonded ComGD-ComGF adducts are cleaved and the two proteins migrate as monomers (Fig. 6). In conclusion, by accurately capturing ComGD-ComGF interactions within the pilus, these results validate the model of the filament tip, especially in the area important for DNA binding.

**Fig. 6.**
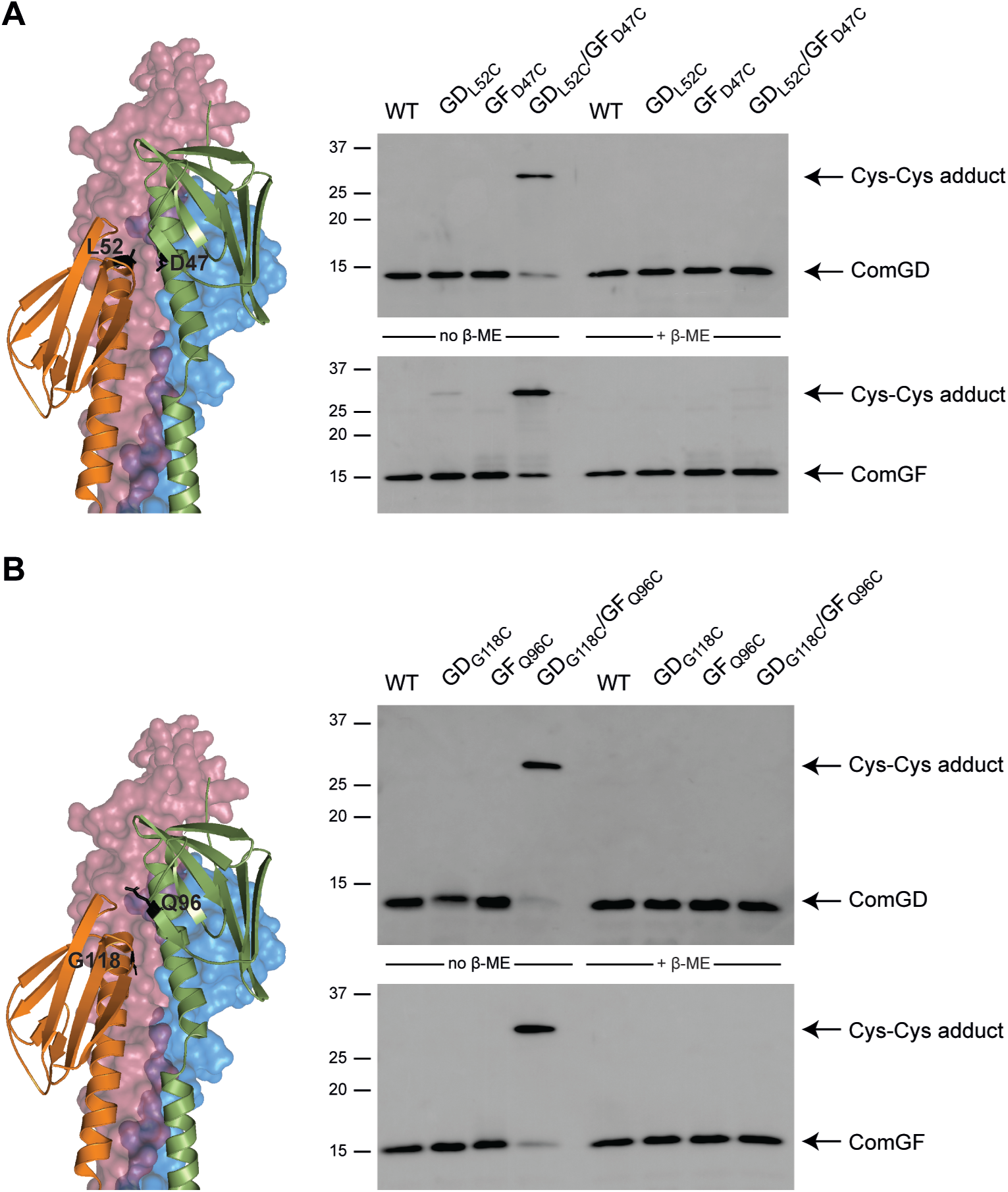
Probing the interface between ComGD and ComGF by Cys crosslinking. We constructed *S. sanguinis* single and double mutants in a strain that constitutively expresses T4dP (17), with Cys substitutions in two different pairs of residues **A**) ComGD_L52C_/ComGF_D47C_ and **B**) ComGD_G118C_/ComGF_Q96C_. Disulfide crosslinking was tested in the presence of 4-DPS oxidizer (32) and disulfide-bonded ComGD-ComGF adducts were detected by immunoblotting on pilus purifications. The adducts were not detected in presence of β-ME reducing agent.

Taken together, these results show that T4dP use an interface between two minor pilins – ComGD and ComGF – present within a tip-located complex of four subunits to promote DNA capture during transformation.

### Competent monoderms use a novel and conserved mode of DNA-binding by two minor subunits of T4P to promote DNA capture during transformation

Using the ability of the recently introduced AlphaFold 3 to predict the joint structure of complexes of proteins with a variety of ligands such as ions, small molecules, and nucleic acids (25), we predicted the structure of *S. sanguinis* T4dP interacting with DNA. Strikingly, as seen in Fig. 7A, the model (only the tip of the pilus is shown) corroborates and further strengthens the findings reported above. The DNA clearly interacts only with one side of the tip-located complex of minor pilins – see second and fourth panel in Fig. 7A – the one formed by the ComGD-ComGF interface. Furthermore, the electropositive residues in ComGD and ComGF identified as important for DNA capture – highlighted in black in Fig. 7A – are positioned at the interface between the two proteins and DNA. Next, we more precisely defined the DNA-binding site used by T4dP to capture DNA (Fig. 7B) by analyzing the complex using PISA (33) (Supplemental Dataset 1) and DNAproDB (34) (Fig. S5). This gave a detailed view of how the two pilins together bind DNA, highlighting a novel mode of DNA-binding (Fig. 7B). It also defined more precisely the areas in ComGD and ComGF that are important for the interaction with the DNA (Fig. 7B), with important residues highlighted in black in the left panel of Fig. 7B. This is strengthened by the confirmation that most of the residues we identified as important in our phenotypic analysis – K_101_, K_123_ in ComGD, and R_73_, K_81_, R_93_ in ComGF – are identified by PISA and DNAproDB.

**Fig. 7.**
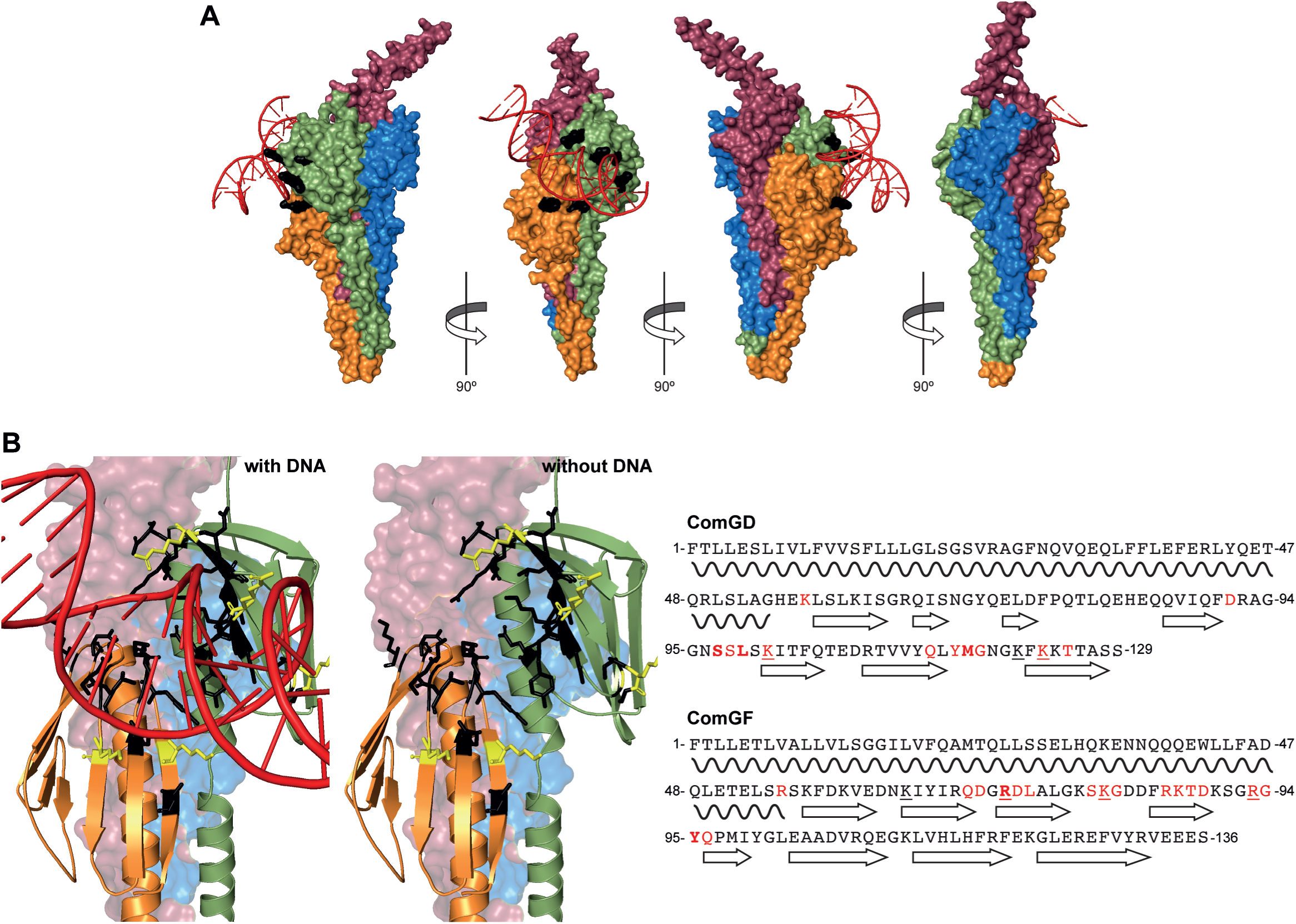
A structural model of the DNA/tip complex in *S. sanguinis* confirms that DNA binds to the ComGD and ComGF interface. **A**) Structural model, predicted by AlphaFold 3 (25), of the pilus tip in *S. sanguinis* in complex with DNA. Since DNA uptake shows no sequence specificity in monoderms, we used a 20 bp portion of pUC19 with 50 % GC content. The surface representation clearly shows that DNA interacts only with by the ComGD-ComGF interface on one side of the tip-located complex, involving the electropositive residues in these minor pilins that we identified as important for DNA (highlighted in black). **B**) DNA-binding site as defined by analysis of the DNA/tip complex using PISA (33) and DNAproDB (34). Left panel, the residues involved in DNA-binding according to PISA and DNAproDB highlighted in black on a cartoon representation, or in yellow when they were also identified in our functional analysis. Right panel, summary of the residues involved in DNA-binding identified in the different analyses, displayed on the sequence of mature ComGD and ComGF. Bold, identified by DNAproDB. Red, identified by PISA. Underlined, identified in our functional analysis. Relevant structural features (α-helix or β-strand) are shown under the sequences.

Finally, we determined whether this novel mode of DNA-binding is conserved in monoderms where T4dP are widespread (9). We generated structural models of tip-DNA complexes in other species where the Com pilus has been extensively studied, *i.e.*, *Bacillus subtilis* and *Streptococcus pneumoniae*. As seen in Fig. 8A, these models are strikingly similar to that from *S. sanguinis*, with DNA interacting specifically with the side of the pilus tip formed by ComGD-ComGF. Bioinformatic analyses of these DNA-protein complex with PISA and DNAproDB show that the DNA-binding sites in these systems are the same than in *S. sanguinis*, with many of the implicated residues ComGD-ComGF being conserved (Fig. 8B). A global multiple sequence alignment (MSA) of the 1,794 ComGD and 2,580 ComGF proteins in InterPro – IPR016785 and IPR016977 entries, respectively – using Clustal Omega (35), show that the residues at the ComGD-ComGF interface important for DNA-binding are broadly conserved (Fig. 8B).

**Fig. 8.**
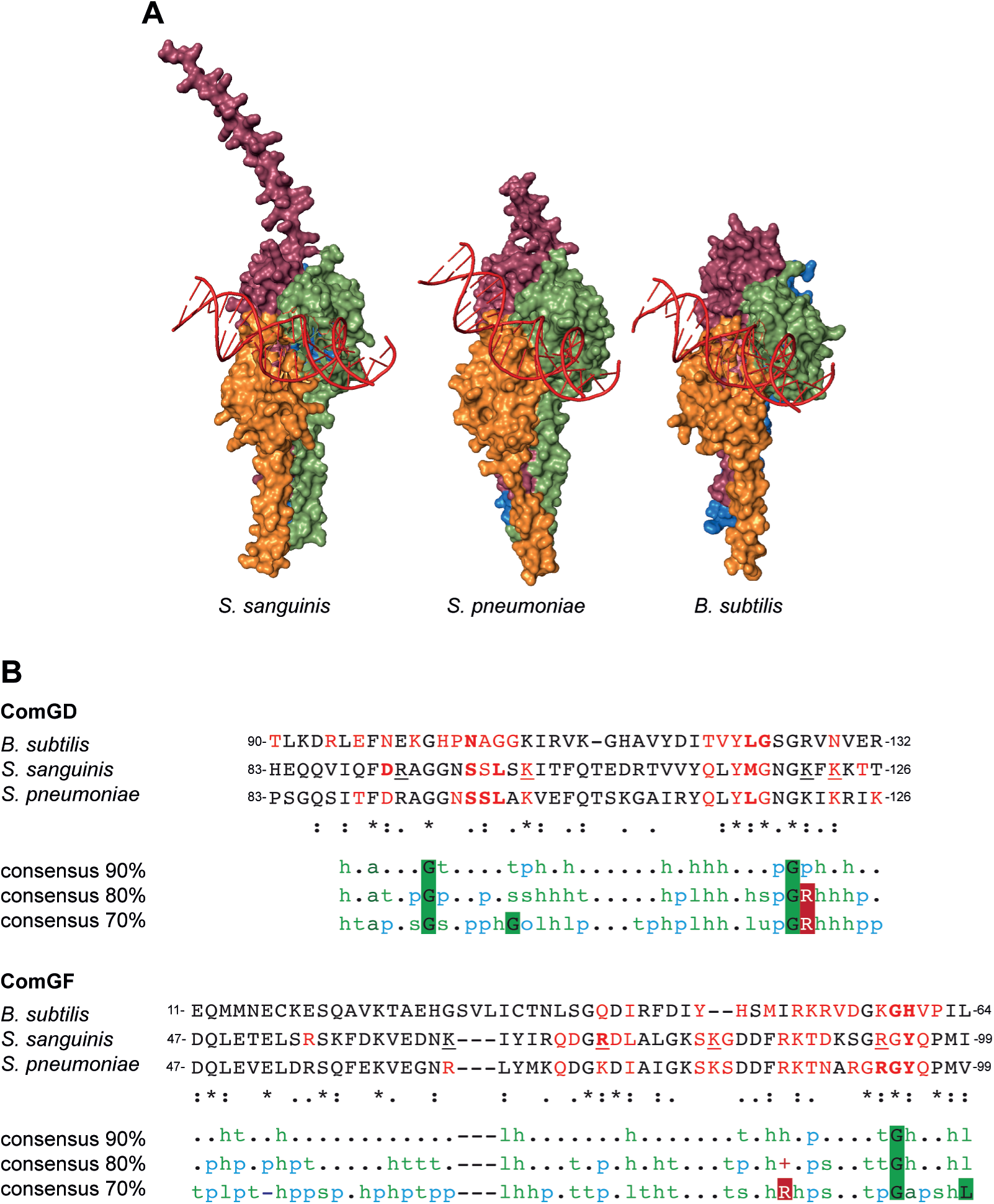
The novel mode of DNA binding we have identified in *S. sanguinis* is broadly conserved in monoderms expressing T4dP. **A**) Structural models, predicted by AlphaFold 3 (25), of the Com pilus tips with bound DNA in *S. pneumoniae* and *B. subtilis* compared to *S. sanguinis*. The DNA always interacts exclusively with the ComGD-ComGF interface. **B**) Comparison of the DNA-binding site as defined by PISA (33) and DNAproDB (34) analyses of the DNA/tip complexes in *B. subtilis*, *S. pneumoniae* and *S. sanguinis*. The sequences were aligned using Clustal Omega (35). The residues involved in DNA-binding by the different analyses were identified as in Fig 7B. The three bottom rows represent the 90, 80 and 70% consensus sequences formatted with MView (44) from MSA computed from the 1,794 ComGD and 2,580 ComGF entries in InterPro. h, hydrophobic (A, C, F, G, H, I, K, L, M, R, T, V, W, Y). t, turn-like (A, C, D, E, G, H, K, N, Q, R, S, T). a, aromatic (F, H, W, Y). p, polar (C, D, E, H, K, N, Q, R, S, T). s, small (A, C, D, G, N, P, S, T, V). o, alcohol (S, T). l, aliphatic (I, L, V). u, tiny (A, G, S). -, negative (D, E). +, positive (H, K, R).

Taken together, these results suggest that T4dP – found in hundreds of species of monoderms – use a highly conserved and previously undescribed mode of DNA-binding implicating two interacting minor pilins located at the pilus tip. This mode of DNA capture is strikingly different than in competent diderms.

## DISCUSSION

Transformation is an important and widespread mechanism of HGT in bacteria, discovered almost 100 years ago (36), which allows competent species to acquire new genetic material (1). This multi-step process involves the capture of free DNA from the extracellular milieu by T4P, that belong to the T4F superfamily of nanomachines ubiquitous in Bacteria and Archaea (8, 9). How T4P bind DNA – the earliest step in DNA capture – remains poorly understood. In a few diderm species, which use T4aP to initiate transformation, minor subunits that directly bind DNA have been identified (11, 12). In monoderms, which use a distinct T4P subtype (T4dP) to promote DNA capture (3), the molecular basis for DNA binding remains unknown. Here, we used *S. sanguinis* – a model competent monoderm species for studying T4F (37) – to define how T4dP interact with free DNA to promote its capture. This led to important findings on key aspects of T4F biology.

The main finding of this study is the unravelling of the mechanism of DNA-binding by T4dP, which appears highly conserved in monoderms. Like in competent diderms – that use T4aP for DNA uptake – where minor pilins such as ComP and FimT displaying intrinsic DNA-binding propensity have been identified (11, 12), we show that minor pilins are involved in DNA capture by T4dP in monoderms. This strongly suggests that in all the species using T4P for DNA capture – virtually all known competent bacteria except *Helicobacter* – specific pilin subunits will act as DNA receptors. However, the similarities end there and the modes of DNA-binding uncovered so far are unexpectedly different. First, in contrast to diderms where DNA-binding is mediated by single pilins such as ComP and FimT (11, 12), monoderms use the interface between two different minor pilins (ComGD and ComGF). Both subunits contribute to DNA-binding and it is only by altering residues in both ComGD and ComGF, which constitute together the DNA-binding site, that competence can be abolished. Although this remains purely speculative, it is likely that this mechanism has evolved by involving at first one of the two pilins in DNA-binding, which was then strengthened by co-opting the second one. Second, ComGD and ComGF are part of a tip-located complex of four pilins required for Com pilus biogenesis, while ComP and FimT are dispensable for piliation and most likely randomly distributed along T4aP (11, 12). Indeed, in the latter filaments, the tip is already occupied by a large adhesin (PilC/PilY1) interacting with a complex of four broadly conserved minor pilins (24). This suggests that different T4P might have evolved the capacity to bind DNA independently during their evolutionary history. Third, although surface-exposed residues always appear to be important, the different pilins playing a role in DNA capture appear to interact with DNA using very different binding sites. This makes it likely that other DNA-binding mechanisms remain to be discovered, notably in bacteria that use T4cP for DNA capture such as *Micrococcus luteus* (38) where no homologs of any of the above DNA-binding pilins are found.

The finding that the complex of four minor pilins at the tip T4dP plays a direct role in their main biological function is of general significance for T4F, because of the analogy with tip-located complexes of four minor pilins in other widespread T4F such as T4aP (24) and type 2 secretion systems (T2SS) (23). This reveals that a set of four minor pilins in different T4F – important in filament biogenesis and likely to have a common evolutionary origin – has been functionalized to play a key role in very different properties. These four tip-located pilins (1) interact and present the PilC/PilY1 adhesin in T4aP (24) that promotes adhesion to a wide variety of surfaces, (2) interact with protein effectors in T2SS (39) to promote their secretion in the extracellular milieu, and (3) bind free DNA to promote its capture by T4dP. Critically, DNA binding by T4dP does not use the most salient structural feature in the tip complex, *i.e.*, the unstructured tail in ComGG, whose role thus remains mysterious. Therefore, this functionalization is yet another evolutionary strategy used by bacteria – in addition to the design of modular pilins grafted with highly diverse functional modules (40, 41) – that contributes to the exceptional functional versatility of T4F.

In conclusion, by identifying the molecular bases of DNA capture by a T4P subtype found in hundreds of species of monoderms and showing that a tip-located complex of four minor pilins found in multiple T4F is involved, this work has general implications for T4F. Moreover, by illustrating the power of experimental approaches based on the use of highly accurate structural models generated by artificial intelligence programs and further cementing *S. sanguinis* as a T4F model, this study paves the way for future investigations that will further improve our understanding of these fascinating filaments.

## MATERIAL AND METHODS

### Structure modelling and bioinformatics

We used AlphaFold 3 (25) on the AlphaFold server for modelling protein 3D structures. We invariably chose the model that was ranked first according to confidence metrics (1) pLDDT (a per-atom confidence estimate) shown as a color output in the image of the structure using the same coloring as in the AlphaFold Protein Structure Database (AFDB) (42), (2) pTM (predicted template modelling score) and (3) ipTM (interface predicted template modelling score). The quality of the complex models was estimated using the structure assessment tool webserver in Expasy (29). We used PyMOL 3.1 (Schrödinger) for molecular visualization and for generating the structure figures. The interfaces in the complexes between proteins and DNA were analyzed using PISA (33) and DNAproDB (34). Signatures in proteins were identified by scanning against the InterPro database (43) using the InterProScan webserver. (21). MSA were performed using Clustal Omega (35) and reformatted using MView (44), on the EMBL-EBI servers. Sequence logos were generated from MSA (45) using the WebLogo 3 server. The two pairs of residues for the Cys crosslinking experiments were selected by using a combination of (i) sequence conservation visualized on the structures using Consurf (46), (ii) co-evolution as assessed by EVcouplings (47), and (iii) the web-based Disulfide by Design 2.0 tool for disulfide engineering in proteins (48).

### Protein expression and purification in *E. coli*

*E. coli* DH5α was used for cloning. *E. coli* BL21(DE3) was used for protein expression and purification. Strains were grown in liquid or solid lysogeny broth (LB) medium (Difco), containing, when required, 100 µg/ml ampicillin.

For constructing expression plasmids (listed in Table S1), we used standard molecular biology techniques (49). We used pMALX(E) to generate maltose-binding protein (MBP) fusions with *S. sanguinis* ComGC and ComEA encoded by genes codon-optimized for expression in *E. coli* (purchased from GeneArt). The portions of *comGC_SS_* encoding ComGC_23-94_, which was previously characterized structurally (30), or *comEA_SS_* encoding residues ComEA_160-226_, were PCR amplified (primers are listed in Table S2), PCR products were cut with *Eco*RI and *Hin*dIII, purified using a QIAquick PCR purification kit (Qiagen). and cloned in pMALX(E) cut by the same enzymes. The resulting plasmids were verified by sequencing.

For protein purification, 1 L cultures grown at 37°C until OD_600_ 0.6 were induced with 0.3 mM isopropyl 1-thio ß-D-galactopyranoside (Sigma) overnight (O/N) at 16°C. Bacteria were harvested by centrifugation at 12,000 *g* and resuspended in lysis buffer (50 mM Tris-HCl pH 8, 100 mM NaCl) or (50 mM Tris-HCl pH 7.5, 200 mM NaCl, 1 mM EDTA) for MBP-ComEA and MBP-ComGC, respectively. Lysis buffer was supplemented with 20 µg/ml DNase I (Sigma) and EDTA-free protease inhibitor cocktail (Roche). Cells were lysed using a French press. The recombinant proteins were affinity-purified using Poly-Prep chromatography columns (Bio-Rad) loaded with amylose resin (BioLabs) and eluted using lysis buffer containing 10 mM maltose. MBP-ComEA was treated 2 h at 30°C with 30 µg/mL of DNase I and 20 mM MgCl_2_. To remove maltose and DNase, proteins were buffer exchanged in lysis buffer using prepacked disposable PD-10 columns (Cytiva) and then a second affinity-purification was done. The eluted proteins were buffer exchanged in EMSA buffer (25 mM Tris-HCl pH 8, 2.5 mM MgCl2, 50 mM NaCl) and concentrated using Amicon Ultra centrifugal filters with a 10 kDa cut-off (Millipore).

### Testing DNA-binding ability of purified proteins

The DNA-binding ability of purified proteins was tested by performing agarose EMSA were performed essentially as described previously (11). In brief, 120 ng pUC19 plasmid DNA was incubated for 30 min at room temperature with increasing concentrations of purified MBP fusion proteins in 10 μl EMSA buffer. DNA was then separated by gel electrophoresis on 0.8 % agarose (Fisher) in Tris acetate-EDTA buffer, and visualized after ethidium bromide staining.

### Construction of *S. sanguinis* strains and growth conditions

All the *S. sanguinis* strains used in this study (listed in Table S1) are derivatives of the 2908 throat isolate (50). *S. sanguinis* strains were grown at 37°C as described, using Todd Hewitt (TH) broth (Difco). Plates, TH broth with 1.5 % agar, were incubated in anaerobic jars (Oxoid) under anaerobic conditions. Liquid cultures in THTH – TH containing 0.05 % <u>T</u>ween 80 (Merck) to limit bacterial clumping, and 100 mM <u>H</u>EPES (Euromedex) to prevent acidification of the medium – were grown statically under aerobic conditions. Strains, constructed as previously described (17) with minor modifications, were all verified by PCR and sequencing. Genomic DNA was prepared from liquid cultures using the XIT genomic DNA from Gram-positive bacteria kit (G-Biosciences). PCR were done using high-fidelity DNA polymerase (Agilent), with primers listed in Table S2. The unmarked mutants were constructed in one step by transforming *S. sanguinis* in the absence of selective pressure, with a splicing PCR product in which the regions upstream and downstream of the target mutation were amplified using F1/R1 and F2/R2 primers (Table S2). For the 5Q mutant construction, we spliced the region containing mutations in *comGD* and *comGF* using as templates the genomic DNA of *comGD_K101Q/121Q/K123Q_* and *comGF_R73Q/R93Q_*, respectively. For the Cys crosslinking experiments, we constructed *S. sanguinis* single and double Cys mutants in ComGD and ComGF mutants in a 2908 derivative that constitutively expresses Com pili (17). This was done to facilitate immunodetection of ComGD and ComGF.

### Transformation of *S. sanguinis*

Competence in *S. sanguinis* strains was quantified as described previously (17). In brief, after O/N growth, bacteria were diluted in THTH. Induction was performed with 300 ng/ml synthetic competence-stimulating peptide (CSP), and we used 100 ng of a purified PCR product encompassing the *rpsL* gene from Str^R^ 2908, a mutant spontaneously resistant to streptomycin. After bath-sonication, we performed serial dilutions that were spread on plates with and without streptomycin. Frequencies were determined as number of Str^R^ transformants/total CFU.

For non-selective transformation, after O/N growth, bacteria were diluted 10^−7^ in pre-warmed THTH and incubated at 37°C for 2 h. We then induced competence in this culture by adding 300 ng/ml synthetic CSP, took a 330 µl aliquot to which we added 500 ng of transforming DNA. After incubation for 2 h at 37°C, bacteria were bath-sonicated before plating to break bacterial chains, using a Bioruptor (Diagenode) at medium amplitude for 60 s. If colonies were non-clonal, as verified by sequencing, bath-sonication was repeated.

### Assaying piliation in *S. sanguinis*

Com pili were purified as previously described (17). Briefly, liquid cultures grown O/N in THTH were used to inoculate pre-warmed THTH at OD_600_ 0.01 and grown statically until the OD_600_ reached 0.04-0.08. CSP was then added at a final concentration of 300 ng/mL, and induction was performed for 30 min. Bacteria were pelleted by centrifugation for 15 min at 4,149 *g* at 4°C. Pili were sheared after resuspending bacterial pellets in ice-cold pilus buffer by repeated pipetting up and down. Bacteria were then pelleted by two rounds of centrifugation at 4°C for 10 min at 9,220 *g*. Finally, pili were pelleted by ultracentrifugation at 100,000 *g* for 1 h at 4°C. The pellets were resuspended in pilus buffer by pipetting up and down.

Pilus preparations were analyzed by immunodetection of the major pilin ComGC. Immunoblotting was done as described previously (17). In brief, proteins were separated by SDS-PAGE in Tris-glycine buffer (Euromedex). The Precision Plus Protein All Blue Pre-Stained Protein Standards (Bio-Rad) was used as molecular weight marker. Gels were transferred onto Amersham Hybond ECL nitrocellulose membrane (GE Healthcare) and analyzed by immunoblotting using anti-ComGC as primary antibody (at 1/2,500 dilution) and anti-rabbit HRP-conjugated (GE Healthcare) as secondary antibody (at 1/10,000 dilution). Detection was performed using Amersham ECL Prime Western Blotting Detection Reagent (GE Healthcare).

Purified pili in the 5Q mutant were visualized by TEM after negative staining as described elsewhere (17). We used a Tecnai 200 KV electron microscope (Thermo Fisher Scientific) and a Oneview 16 Megapixel camera (Gatan) to acquire images.

### Cys crosslinking experiments

Pilus purification for the Cys crosslinking experiments in strains that constitutively expresses T4dP were done as follows. Liquid cultures grown O/N in THTH were used to inoculate pre-warmed THTH at 10^−8^ and grown statically until OD_600_ reached 0.6. Bacteria were pelleted by centrifugation for 15 min at 4,149 *g* at 4°C. Pili were sheared after re-suspending bacterial pellets in PBS (Sigma) by repeated pipetting up and down. Bacteria were then pelleted by two rounds of centrifugation at 4°C for 10 min at 9,220 *g*. Pili were then incubated with 200 µM of 4-DPS (Sigma) for 30 min on ice. Finally, pili were pelleted by ultracentrifugation at 100,000 *g* for 1 h at 4°C. The pellets were resuspended with PBS by pipetting up and down. Sample were treated with and without β-mercaptoethanol (Aldrich) and analyzed by SDS-PAGE and immunoblotting as described above, using anti-ComGD or anti-ComGF as primary antibodies (at 1/1,000 dilution). Detection was performed using SuperSignal West Atto Ultimate Sensitivity Chemiluminescent Substrate (Thermo Scientific).

## ACKNOWLEDGEMENTS

This work was supported by the Agence Nationale de la Recherche (ANR-21-CE11-0008-01). We thank Artemis Kosta and Hugo Le Guenno (Plateforme de Microscopie, Institut de Microbiologie de la Méditerranée, Marseille) for help with electron microscopy. We are grateful to colleagues from the Laboratoire de Chimie Bactérienne – Emilia Mauriello and Romé Voulhoux – for critical reading of this manuscript.

